# Expression and DNA methylation of 20S proteasome subunits as prognostic and resistance markers in cancer

**DOI:** 10.1101/2023.12.12.571247

**Authors:** Ruba Al-Abdulla, Simone Venz, Ruslan Al-Ali, Martin Wendlandt, Mandy Radefeldt, Elke Krüger

## Abstract

Proteasomes are involved in the maintenance of cellular protein homeostasis and the control of numerous cellular pathways. Single proteasome genes or subunits have been identified as important players in cancer development and progression without considering the proteasome as a multi-subunit protease. We here conduct a comprehensive pan-cancer analysis encompassing transcriptional, epigenetic, mutational landscapes, pathway enrichments, and survival outcomes linked to the 20S proteasome core complex. The impact of proteasome gene expression on patient survival exhibits a cancer-type dependent pattern. Escalated proteasome expression associates with elevated activation of oncogenic pathways, such as DNA repair, MYC- controlled gene networks, MTORC1 signaling, oxidative phosphorylation, as well as metabolic pathways including glycolysis and fatty acid metabolism. Vice versa, potential loss of function variants correlates with improved survival. The TCGA-derived outcomes are further supported by gene expression analysis of THP-1 cells. Our study reframes these subunits as an integrated functional ensemble, rather than separated subunits.

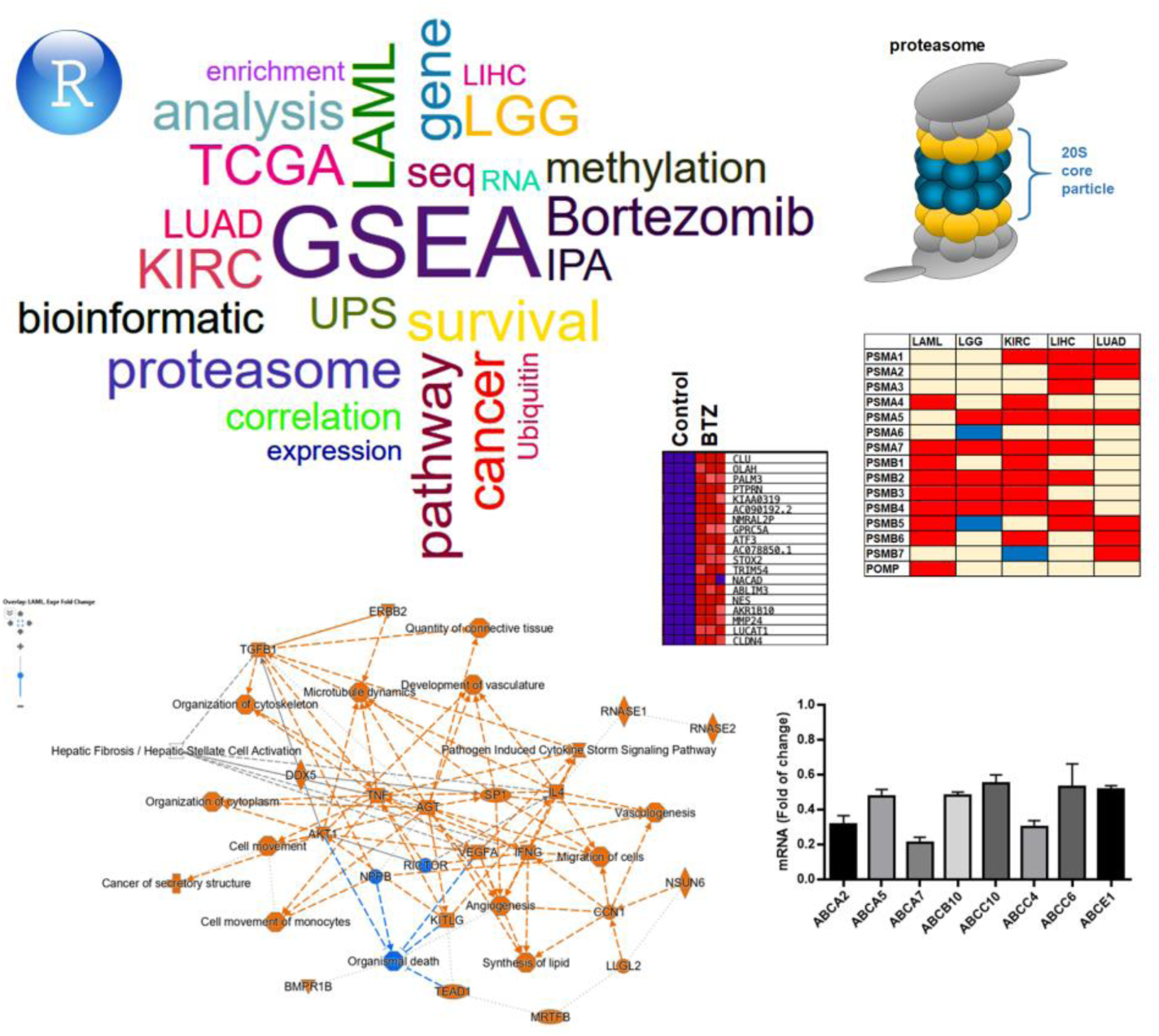

## 1. Introduction

Protein homeostasis plays a decisive role in maintaining cellular function and survival. In both healthy and diseased states, cells meticulously sustain an equilibrium between protein synthesis and degradation through the process of protein quality control [1]. The ubiquitin proteasome system represents the primary intracellular and non-lysosomal mechanism for protein degradation within cells [2]. The proteasome, a molecular machinery with a highly conserved architecture and protease activity, plays a pivotal role in the selective degradation of intracellular proteins, including misfolded proteins, regulatory proteins, and signaling proteins. Thereby, the proteasome controls cellular processes, including immune regulation, cell cycle progression, apoptosis, and DNA repair [3–8]. Protein degradation by the proteasome is initiated by a thioester cascade involving the enzymes of the ubiquitin conjugation machinery E1, E2 and E3, which mediate polyubiquitination of protein substrates as prerequisite for degradation by the proteasome [3]. Polyubiquitinated proteins with lysin 48-linked ubiquitin chains are typically degraded by the 26S proteasome complex consisting of two sub complexes, 20S core particle (CP) and 19S regulatory particle (RP) [9, 10]. The 20S core particle exhibits a barrel-like structure made out of four heptameric rings which are themselves composed of ɑ- and β-subunits. The two outer rings comprise 7 ɑ-subunits encoded by *PSMAs* (*PSMA1-7*). The inner two β-subunits rings are encoded by *PSMBs* (*PSMB1-7*) [7] which contain the catalytically active threonine sites. Incorporation of β1, β2, and β5 into the 20S complex ensures the caspase-, trypsin- and chymotrypsin- like activities, respectively [9]. Beyond the standard proteasome (SP), a spectrum of alternate isoforms, including the immunoproteasome (IP), has garnered significant attention. The IP is not only expressed within immune cells but can also be induced in diverse cell types, spurred by specific cytokine cues. Distinguished by the incorporation of inducible β1i-, β2i-, and β5i-subunits (encoded by *PSMB8-10*), the IP demonstrates an augmented proteolytic potency. [3, 11]. Dysfunction and dysregulation of the proteasome has been linked to a wide range of diseases, including cancer, neurodegenerative disorders, and autoimmune diseases [12–14]. Several studies have reported alterations in the expression of single proteasome subunits in various types of cancer. *PSMA1*, a subunit of the 20S CP, has been found to be overexpressed and considered a prognostic or cancer marker in several tumors, including lung, gastric, breast and colon cancer [15–18]. Indeed, proteasome α-subunits have been associated with the development and progression of several types of cancers, including hepatocellular carcinoma, breast cancer, lung cancer, and multiple myeloma [19–23]. Certain β-subunits were as well identified as therapeutic targets in solid tumors such as hepatocellular carcinoma or kidney cancer [24, 25]. However, single subunits cannot function in selective degradation of target proteins. Therefore, there is an urgent need to evaluate the impact of proteasome expression and activity in cancer as an enzymatic unit, rather than the evaluation of the role of separated subunits. Proteasome expression in cancer is associated with oncogenic signaling pathways, such as the PI3K/Akt/mTOR, NF-κB, Cyclin D1 signaling, interferon signaling and Wnt-β catenin pathways [26–30].

In addition, elevated proteasome expression has been associated with the development, progression, and therapeutic resistance of tumours. Therefore, targeting proteasomes via pharmacological inhibition potentiates the anti-cancer efficacy of other chemotherapeutic drugs [27]. Given the dysregulation of proteasome expression in cancer, SP and IP inhibitors have emerged as a promising class of cancer therapeutics [12]. The first proteasome inhibitor, bortezomib (BTZ), was approved by the FDA. This medication has shown significant efficacy in the treatment of multiple myeloma, mantle cell lymphoma, and several types of solid tumors [31, 32]. Several other proteasome inhibitors have been developed and are currently being investigated as cancer therapeutics [33].

In the current study we aimed to answer two questions; i) In which type of cancers does the proteasome 20S expression possess a prognostic value, and ii) what are the oncogenic pathways by which high expression of proteasome have a worse impact on patient survival.

## 2. Methods

### 2.1. Survival analysis

To investigate the impact of the transcriptome levels of proteasome 20S subunits, namely *PSMA1* (NM_002786.3), *PSMA2* (NM_002787.4), *PSMA3* (NM_002788.4), *PSMA4* (NM_002789.4), *PSMA5* (NM_002790.3), *PSMA6* (NM_002791.1), *PSMA7* (NM_002792.3), *PSMB1* (NM_002793.3), *PSMB2* (NM_002794.4), *PSMB3* (NM_002795.2), *PSMB4* (NM_002796.2), *PSMB5* (NM_002797.3), *PSMB6* (NM_002798.2), *PSMB7* (NM_002799.3) as well as *POMP* (NM_015932.5), on patients overall survival, we performed a survival analysis using Kaplan-Meier (K-M) survival curves obtained from the TCGA portal (http://tcgaportal.org/index.html). The median expression level of each gene was used as a cutoff value. Our findings were confirmed using the K-M-plotter tool (https://kmplot.com/analysis/). The results are presented in a color-coded table, where red indicates that elevated gene expression is associated with worse prognosis, blue indicates that high expression is associated with better survival, and pale yellow indicates that the association was not significant. We considered results statistically significant if the Log-rank p-value was less than 0.05.

### 2.2. DNA methylation and overall survival of patients in 5 types of cancer

In this study, we investigated the impact of DNA methylation on the overall survival of patients in five types of cancer (Kidney renal clear cell carcinoma (KIRC), Acute Myeloid Leukemia (LAML), Low-Grade Gliomas (LGG), Liver Hepatocellular Carcinoma (LIHC) and Lung adenocarcinoma (LUAD)) by analyzing the DNA methylation status of 20S proteasome subunits. To achieve this aim, survival curves for each CpG probe located in these genes were obtained from the publicly available MethSurv database (https://biit.cs.ut.ee/methsurv/). DNA methylation values were represented as beta values, which ranged from 0 to 1, and the median was used as a cut-off value [34]. Results were considered statistically significant if consistent results were obtained with a Log-rank p-value less than 0.05. To present the results, we used a color-coded table, where blue indicates that high levels of DNA methylation are associated with better patient survival, and red indicates that high levels of DNA methylation are associated with worse survival. Data of DNA methylation were downloaded, and the correlation between DNA methylation and RNA expression was determined using a R-based code for Spearman factor.

### 2.3. Variant interpretation of TCGA somatic sequencing data

Somatic mutations associated with proteasome 20S subunit genes, were extracted from the c-bioportal (TCGA/Pancancer Atlas dataset), which included data from 5 different types of cancer. Variants were pre-selected using the aggregator and impact analysis tool VarSome [35] based on factors such as predicted consequence, structural and functional impact, and phylogenetic conservation. Subsequently, the resulting variants were then manually classified following the updated American College of Medical Genetics and Genomics (ACMG) criteria [36]. Nomenclature for genetic information adheres to the recommendations provided by the Human Genome Variation Society (HGVS) [37]. For structural analysis, mature 20S subunit protein sequences containing a substitution of interest were subjected to folding using AlphaFold. Subsequently, these folded structures were compared with their corresponding reference structures in the context of the complete 20S proteasome core particle obtained from PDB ID 5LF3, using ChimeraX version 1.6.1. This comparison also allowed for the visualization of hydrogen bonds, electrostatic potential and hydrophobicity.

### 2.4. Pathway analysis

The pathway analysis was determined using the gene expression files downloaded from c-bioportal-TCGA/Pancancer dataset. Data was then stratified based on the expression of both *PSMAs* (1-7), *PSMBs* (1-7), and *POMP*. High expression was defined as the expression of at least 10 out of the 15 genes above the median, otherwise it was classified as sample with low expression. This stratification is carried out using Anaconda-Jupyter python code, and followed by a pathway analysis using the gene set enrichment analysis (GSEA) [38, 39]. We applied the next parameters (1000 repeat, Reactome and hallmark pathway studies) using a cutoff p-value of 0.05 with FDR value of 0.25. For the seek of simplicity we only showed the results of the hallmark platform. Confirmation of the results was carried with a second strategy of stratification using the mean of the 15 genes, and apply the median of this mean as a cut off value afterwards (Data is provided upon request).

Gene ontology, functional analyses as well as pathway network analyses were performed with the expression data using the Ingenuity Pathways Analysis tool (IPA, QIAGEN Bioinformatics, Redwood, CA, USA; https://www.qiagenbioinformatics.com/products/ingenuity-pathway-analysis/). The z-Score algorithm was employed to forecast the direction of alteration in the present activity. IPA’s z-score is a measure that forecasts the activation or inhibition of a pathway or gene. A negative z-value (blue) suggests that the pathway is mostly inhibited, while a positive z-value (orange) indicates that the pathway is predominantly activated. According to company descriptions (https://qiagen.my.salesforce-sites.com/KnowledgeBase/KnowledgeNavigatorPage?id=kA41i000000L5pACAS&categoryN <u>ame=IPA</u>) the canonical pathways analysis exhibits the most noteworthy pathways in the complete dataset, with significance determined by the right-tailed Fisher’s Exact T-test, indicating the likelihood of association between molecules from the dataset and the canonical pathway occurring randomly. The data from the analysis were visualized as a bubble chart, in which the y-axis shows the pathway categories, and the x-axis shows their z-score. The pathway bubbles are colored based on their z-score and the bubble size is proportional to the number of genes from the dataset that overlap with each pathway.

### 2.5. Cell Culture and RNA-seq analysis

THP-1 (RRID: CVCL_0006) cells were cultivated in RPMI medium supplemented with FCS 10% and Penicilin/ Streptomycin and incubated at 37°C and CO_2_ 5%. Cells were treated with Bortezomib at a non-toxic dose (10 nM) for 16 hrs. After collection, RNA was extracted and sequenced at Centogene GmbH-Rostock. RNA quality was assessed with the RNA Screentape on the Agilent Tapestation (Agilent, Santa Clara, CA).

RNA sequencing: RNA libraries were prepared using the Illumina stranded mRNA kit (Illumina, San Diego, CA). Quality Control (QC) of the libraries was conducted with the DNA 1000 Screentape on an Agilent Tapestation. Samples were sequenced on an Illumina NextSeq500 using the NextSeq 500/550 High Output v2.5 reagents and the 75 bp paired-end protocol. RNA-seq raw data was aligned using two-pass mode with STAR v.2.7.6a against hg19 [40]. The read groups are fixed, and the duplicates marked using Picard tools v.2.23.8 (http://broadinstitute.github.io/picard). Counting the reads was performed by feature Counts/subread v.2.0 [41]. Pathway enrichment analysis was performed with Gene Set Enrichment Analysis (GSEA) Gene differential analysis [38]. Gene differential analysis was additionally confirmed with DESeq2 [42] and Toppgene [43].

### 2.6. Correlation study

The correlation between gene expression of 20S proteasome subunits and specific genes was carried out using GEPIA2 [44]. Pearson factor was used to estimate the correlation. Gene signature of the proteasome 20S subunits was created using the 15 genes included in this study. Gene signature of ABC transporters was created using 24 genes of the ABC family of transporters (*ABCA1, ABCA2, ABCA3, ABCA5, ABCA7, ABCB10, ABCB4, ABCB6, ABCB7, ABCB8, ABCB9, ABCC1, ABCC10, ABCC4, ABCC5, ABCC6, ABCD1, ABCD3, ABCD4, ABCE1, ABCF1, ABCF2, ABCF3,* and *ABCG1*). These genes were chosen due to the fact that they are expressed in THP-1cells which facilitate the validation of the obtained data. A correlation was considered very week if R value is below 0.3, week if R value between 0.3 and 0.7. A strong correlation is observed when R value is above 0.7. The correlation between gene expression and DNA methylation probes was carried out using data obtained from MethSurv. Spearman factor was calculated for each CpG probe and the corresponding gene. Results were considered significant if p value is less than 0.05. Original data tables are provided in the supplementary data.

## 3. Results

### 3.1. Correlation between proteasome expression and survival of patients with cancer

First, we aimed to assess the prognostic value of proteasome subunit expression in different cancer types available in the cancer genome atlas (TCGA) database by using Kaplan-Meier survival curves. Our findings revealed a distinct cancer type-dependent pattern of association between the genetic expression of proteasome subunits and patient survival (Figure 1). Supplementary Table S1 provides the abbreviations for these cancer types along with the number of patient samples analyzed in each category. Additionally, the abbreviations for proteasome genes and their corresponding proteins are presented in Supplementary Table S2.

**Figure 1:**
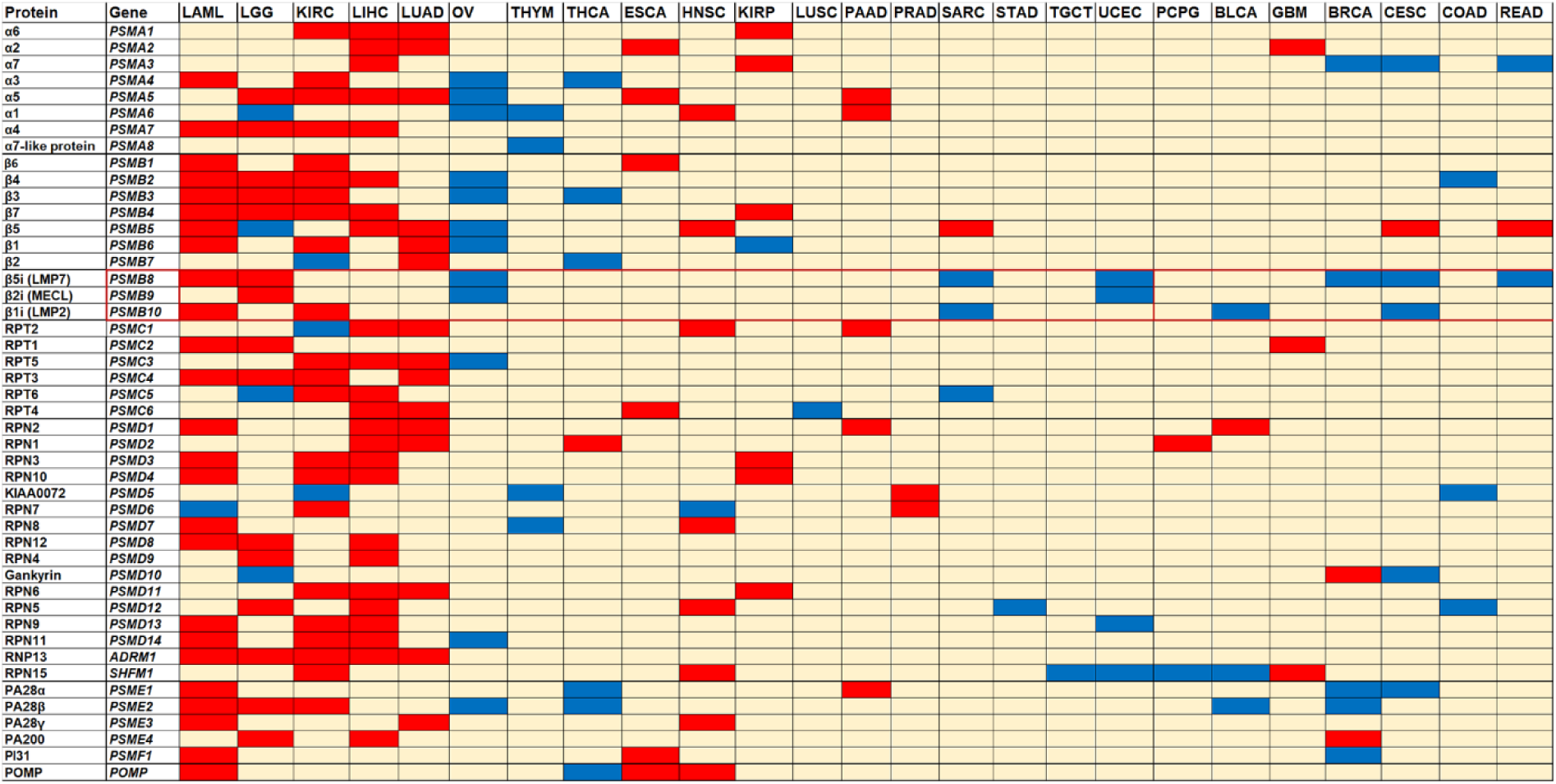
Association between proteasome subunit expression and overall survival in patients with different types of cancers. Kaplan Meyer survival curve for each gene is obtained in each type of cancer from TCGAPORTAL (Abbreviation and sample number are provided in Supplementary Table S1). Median is used as a cutoff value. Difference is considered significant if p ≤ 0.05. Results for association of high expression and survival are presented in color code: Red - worse overall survival; Blue - better overall survival; pale Yellow - no significant association.

In the case of acute myeloid leukemia (LAML) and hepatocellular carcinoma (LIHC), high expression of proteasome subunits was predominantly associated with poorer survival outcomes. Conversely, the pattern observed in other tissues such as the brain, kidney, and lung varied depending on the cancer type. Notably, proteasome subunits expression did not significantly impact patient survival in glioblastoma multiforme (GBM), kidney renal papillary cell carcinoma (KIRP), and lung squamous cell carcinoma (LUSC). However, for low grade glioma (LGG), kidney renal cell carcinoma (KIRC), and lung adenocarcinoma (LUAD), high expression of proteasome subunits genes was linked to worse patient survival. On the other hand, ovarian serous cystadenocarcinoma (OV) and, to a lesser extent, thyroid carcinoma (THCA) demonstrated a favorable prognostic value of proteasome subunits expression. In these cases, elevated expression of proteasome subunits was associated with improved overall survival. Specifically, the expression levels of proteasome subunits in adrenal cancer, breast cancer, esophageal cancer, stomach adenocarcinoma, colon cancer, pancreatic cancer, prostate cancer, thymus cancer, testicular cancer, cervical cancer, and uterine cancer did not exhibit a significant association with altered patient survival. The IP subunits (*PSMB8-10*) exhibited a distinct pattern compared to the standard proteasome. While worse prognosis was primarily observed in LAML and LGG, enhanced patient survival was noted in OV, uterine corpus endometrial carcinoma (USEC), cervical squamous cell carcinoma and endocervical adenocarcinoma (CESC), and sarcoma (SARC) (Figure 1). For the sake of simplicity, we focused our further analysis on the genes encoding the SP 20S subunit, which encompass *PSMA1-7, PSMB1-7*, and the proteasome assembly protein *POMP*. The most relevant results where high proteasome 20S expression is associated with worse survival was observed in LAML, LGG, LIHC, LUAD and KIRC. Therefore, we restricted our further analysis to these types of tumors.

### 3.2 Gene expression, DNA methylation of proteasome 20S subunit genes and overall survival of patients in cancer

It is well established that DNA methylation is involved in the regulation of gene expression and such a regulation is disturbed in cancer [45]. In the current study, we aimed to investigate if hypomethylation of the proteasome 20S subunits is associated with worse overall survival of cancer patients. Therefore, further analysis was conducted using TCGA data to evaluate the association between DNA methylation of the 20S proteasome subunits and overall survival of patients. We evaluated gene expression (obtained from c-bioportal) and DNA methylation (obtained from UALCAN-TCGA) in LIHC, LAML, LGG, LUAD and KIRC. The expression of proteasome subunits was found to be abundant in cancer (Figure 2A). The most elevated levels of expression are seen in LIHC which was associated with lower methylation levels (Figure 2B). We observed higher expression of *PSMB1, PSMB3* and *PSMB4* in tumor samples when compared to other subunits (Figure 2A), Moreover, elevated expression in tumor was observed when compared with non-tumor surrounding tissue in LIHC, LUAD, and KIRC for almost all proteasome subunits. The highest levels of DNA methylation were observed in *PSMA4* and *PSMB5* (Figure 2B). The levels of DNA methylation of *PSMB2* were higher in LUAD, KIRC and LGG, and lower DNA methylation of this subunits was observed in LAML, and LIHC (Figure 2B). Lower levels of DNA methylation of the proteasome 20S subunits were observed in tumor samples when compared to non-tumor surrounding tissue in LIHC, and LUAD. However, in KIRC, a slight elevated DNA methylation was observed in tumor samples when compared with non-tumor tissue. Indeed, our results indicates that hypomethylation of the proteasome 20S subunits is associated with worse survival in LIHC, LGG and LAML.

**Figure 2:**
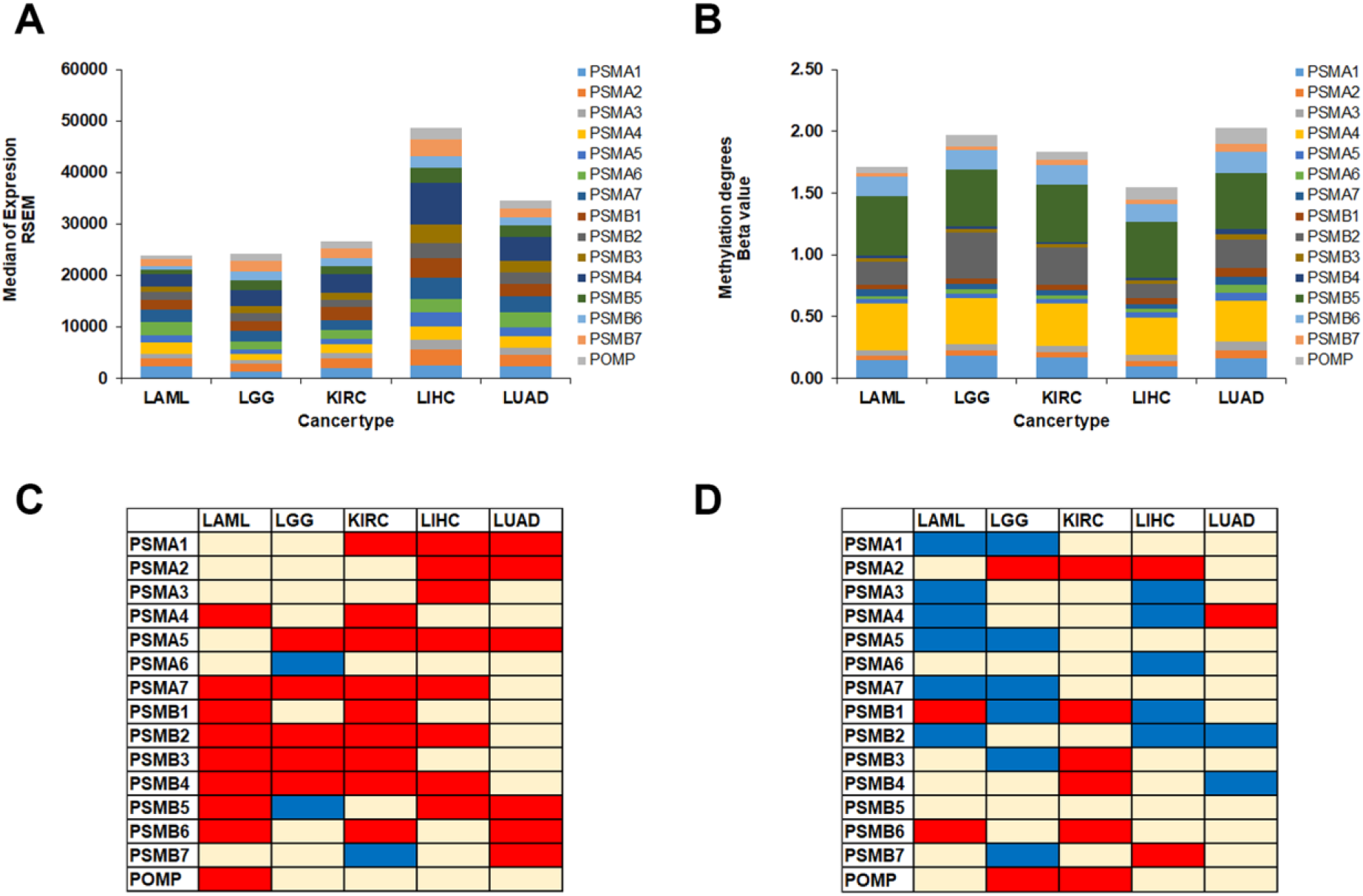
Gene expression, DNA methylation of proteasome 20S subunit genes and patient survival in cancer for LAML, LGG, KIRC, LIHC and LUAD. (A) Gene expression of proteasome 20S subunit genes calculated as the median of RSEM in TCGA samples; (B) DNA methylation of the abovementioned genes represented as beta value for the probes of the promoter obtained from UALCAN database; (C-D) colour-based table of Kaplan Meyer survival of cancer patients based on the expression of proteasome subunits. Association of high gene expression with better survival (coloured in blue) and worse survival (coloured in red), and absence of significant differences was coloured in pale yellow. Median was used as a cut off value, and differences were considered significant when p ≤ 0.05.

The correlation between DNA methylation and gene expression of each subunit was then carried out using spearman factor. In LGG (Figure 3B) a significant negative correlation is observed between the expression of *PSMA5* and *PSMB2* and the methylation degree of several probes that are located in these genes which indicates that DNA methylation could be a mechanism by which the expression of these genes is regulated. Similar results were obtained for the DNA methylation of *PSMA5* and *PSMB4* in LIHC (Figure 3C). No marked negative correlation was observed between the expression of proteasome 20S subunit genes and DNA methylation at the corresponding probes in LAML, KIRC and LUAD as observed in Figure 3A, 3D and 3E respectively.

**Figure 3:**
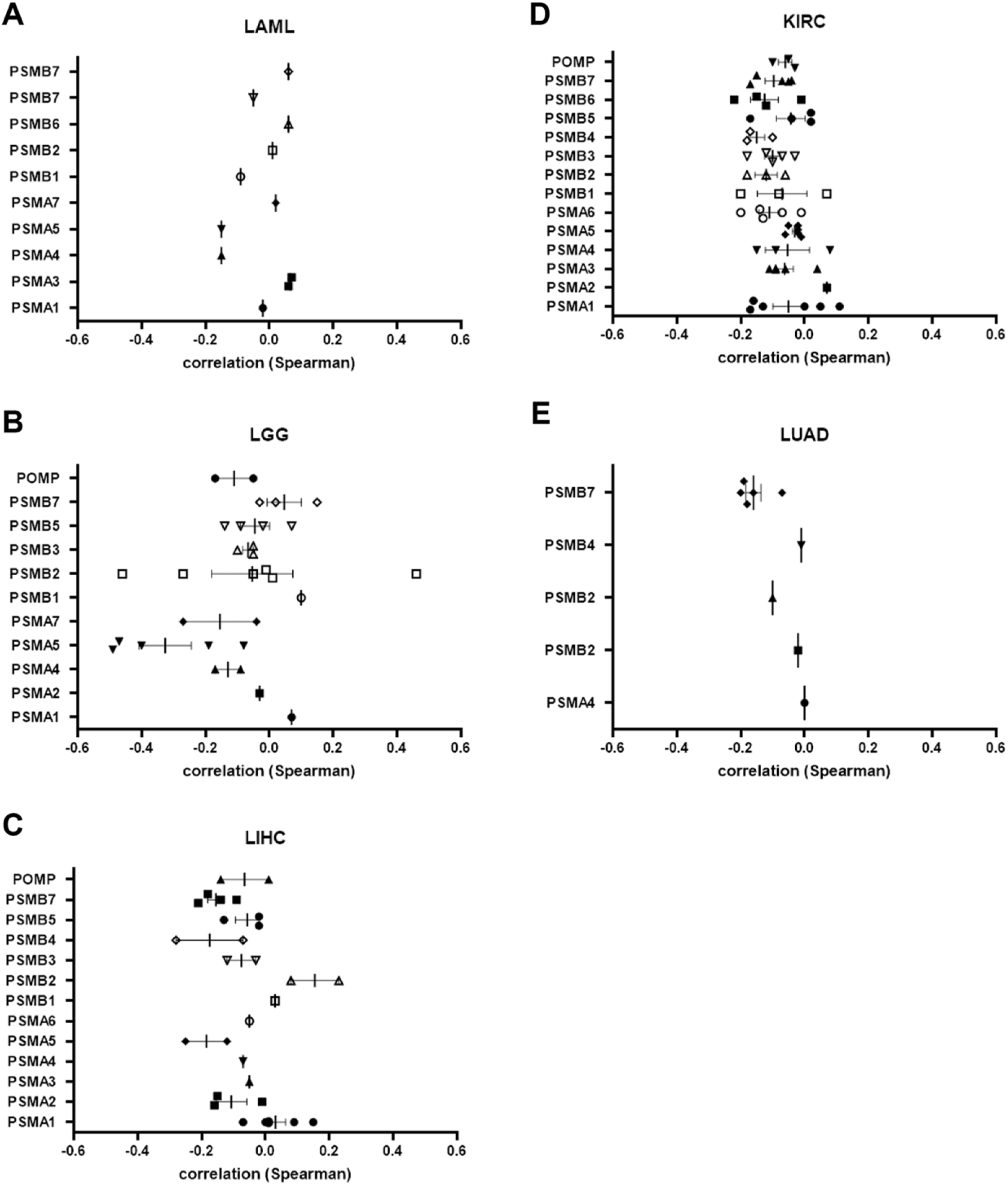
Correlation between DNA methylation and expression of proteasome 20S subunit genes. Box blot of Spearman correlation factor of CpGs probes and gene expression of each gene in (A) LAML; (B) LGG; (C) LIHC; (D) KIRC; (E) LUAD. DNA methylation data (β-value for each probe) and gene expression data (RSEM) were obtained from metsurv TCGA pan-cancer data. Each point represents a correlation of specific DNA methylation probe with gene expression of the corresponding gene.

### 3.3 Variant identification of TCGA somatic sequencing data

Elevated gene expression, and DNA hypomethylation of the 20S proteasome subunits was found to be associated with worse overall survival in patients with LAML, LIHC, KIRC, LGG and LUAD. To study if loss of function of the enzymatic activity of the proteasome is in turn associated with improved survival, we have studied a specific LGG case in which multi-mutations in different proteasome subunits were detected. A total of 112 variants were identified by extracting data from the TCGA dataset for the 15 proteasome-associated genes and narrowed down to 16 variants of interest after pre-selection process using VarSome and ACMG criteria. Among them, we discovered 6 likely pathogenic variants (ACMG Class 4) and 10 variants of uncertain significance (ACMG Class 3) that exhibited a stronger indication of a potential functional impact. Specifically, these variants consisted of 3 frameshift variants, 4 canonical splice variants, and 9 missense variants, as summarized in Supplementary Table S3. All identified variants were absent in normal tissue samples and occurred at frequencies ranging from 0.06 to 0.69. One sample (TCGA-DU-6392) with the variant p.Arg86His in β7/PSMB4 stood out in particular. This patient, diagnosed with LGG, demonstrated an exceptionally extended survival period, surpassing the median survival by almost 10-fold (Figure 4A). Upon further investigation, it became evident that this particular sample harbored the highest number of 20S subunit variants and 6 of them being located in the SP core particle (Figure 4B-H): α6/PSMA1:p.Asp218Tyr, α2/PSMA2:p.Gln96Arg, α7/PSMA3:p.Glu204Lys, α3/PSMA4:p.Ala39Val, β4/PSMB2:p.Lys37Asn, β7/PSMB4:p.Arg86His. All variants were exclusive to the patient’s tumor samples and not present in the normal tissue. According to the predictions, the variants may have varying degrees of considerable impact, notably, the β7/PSMB4 and the α7/PSMA3 variant, with a somatic frequency of 0.44 and 0.30, respectively, were predicted to exert the most profound consequences among them. In the β7/PSMB4 reference structure, p.Arg86 is anticipated to interact with an upstream aspartate at position 70, possibly in conjunction with an arginine at position 227 (not shown), forming stabilizing hydrogen bonds. However, in TCGA-DU-6392, these hydrogen bonds are disrupted by the histidine substitution at position 86, likely resulting in the destabilization of this region (Figure 4H). This region serves as a contact area with the adjacent β1/PSMB6 subunit, apart from the β7/PSMB4 C-terminal interaction with the β1/PSMB6 catalytic domain of the opposite β-ring.

**Figure 4:**
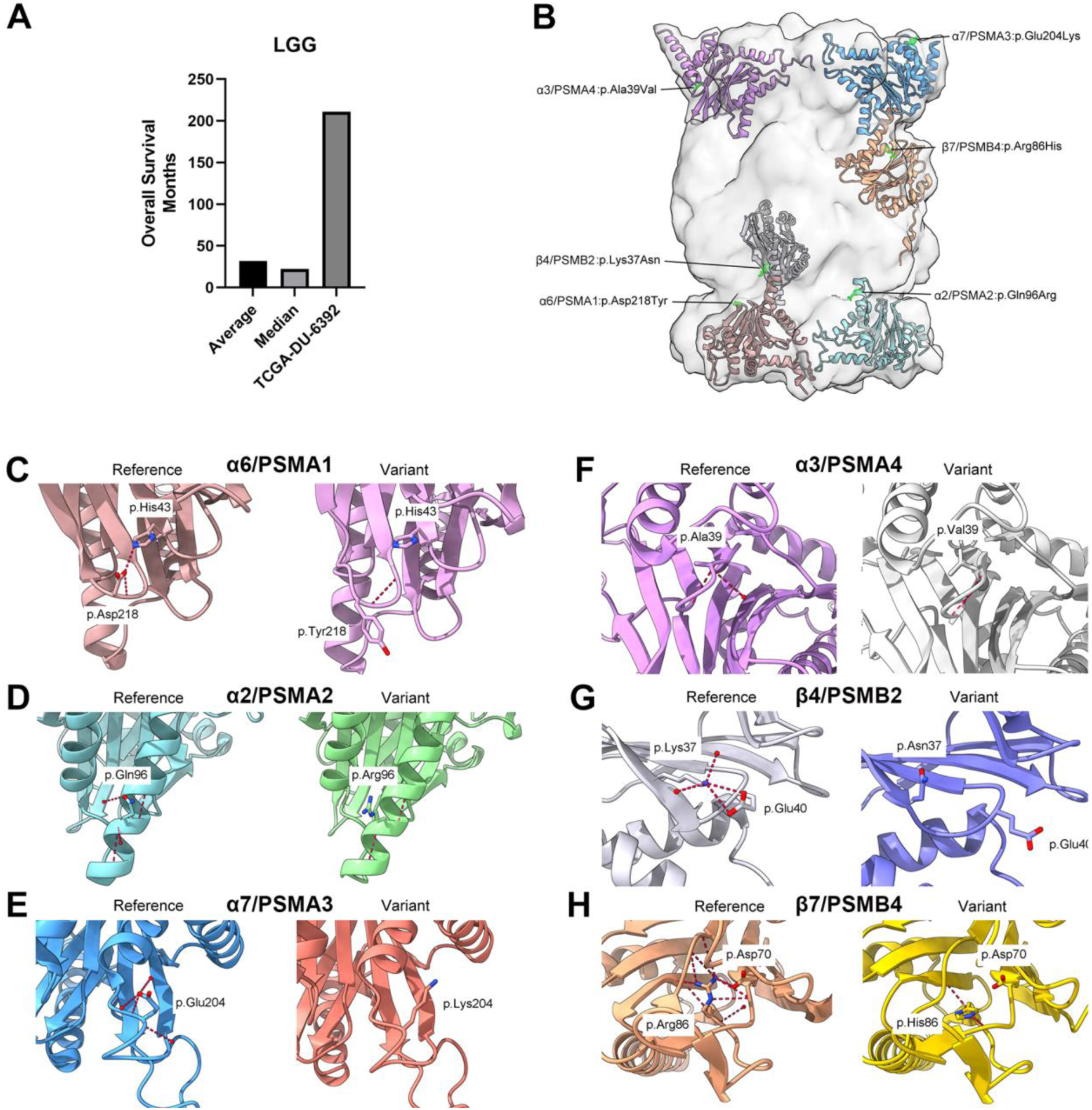
Survival and structural representation of the variants found in sample TCGA-DU-6392. (A) Survival of TCGA-DU-6392 compared to the average and median LGG survival in months. (B) Location of the six standard core particle variants within each respective subunit. (C-H) Comparison between each subunit’s reference structure (left) and the corresponding folded variant structure (right) of this sample. Red dotted lines represent hydrogen bonds, and residues that are putatively interacting with the identified variants are also illustrated.

The glutamate at position 204 in α7/PSMA3 is a component of a turn that establishes contact with the 19S particle, particularly the Rpt3/PSMC4 subunit. The significant change from the negatively charged glutamate to the positively charged lysine at this position in the sample is expected to induce destabilization in this region and disrupt the interaction with the 19S particle (Supplementary Figure S1C). These results indicate a possible association between prolonged survival and reduced proteasome function.

### 3.4 Pathway enrichment in patients with high and low 20S proteasome subunit expression

To estimate the functional role of 20S proteasome subunits function in cancer, we carried out a pathway enrichment analysis in the 5 types of cancer included in the current study. Patients samples were stratified based on the expression level of the median of proteasome subunits (HIGH vs LOW). Pathway enrichment analysis is then carried out using the GSEA software applying using the hallmark platform (Figure 5). Results were presented as a blot bars in red (enriched in the high expression group) and blue (enriched in the low expression group). We have found elevated expression of several pathways in all studied types of cancer (Figure 5A-E). These pathways include DNA repair, MYC targets, MTORC1 signaling, oxidative phosphorylation, reactive oxygen species, and metabolic pathways such as glycolysis and fatty acid metabolism. In the low expression group, except for LAML, a significant enrichment is observed in the Hedgehog signaling pointing to dysregulated stem cell differentiation [46]. While the upstream part of the UV response was enriched in the high expression group, the downstream part of this pathway was enriched in the low expression group. Enrichment of TGF-β pathway was observed in the low expression group of LIHC and KIRC (Figure 5G and I respectively). To validate the results, the analysis was repeated using the mean of each sample for the 15 genes with median cut off value for stratifications. With this method we obtained similar data. Elevated expression of ABC transporters is known to be associated with elevated resistance in cancer [47]. When Reactome platform was used to carry out this analysis, a significant enrichment in ABC transporter, stem cells markers and RNA-binding proteins were observed in the group with high proteasome expression.

**Figure 5:**
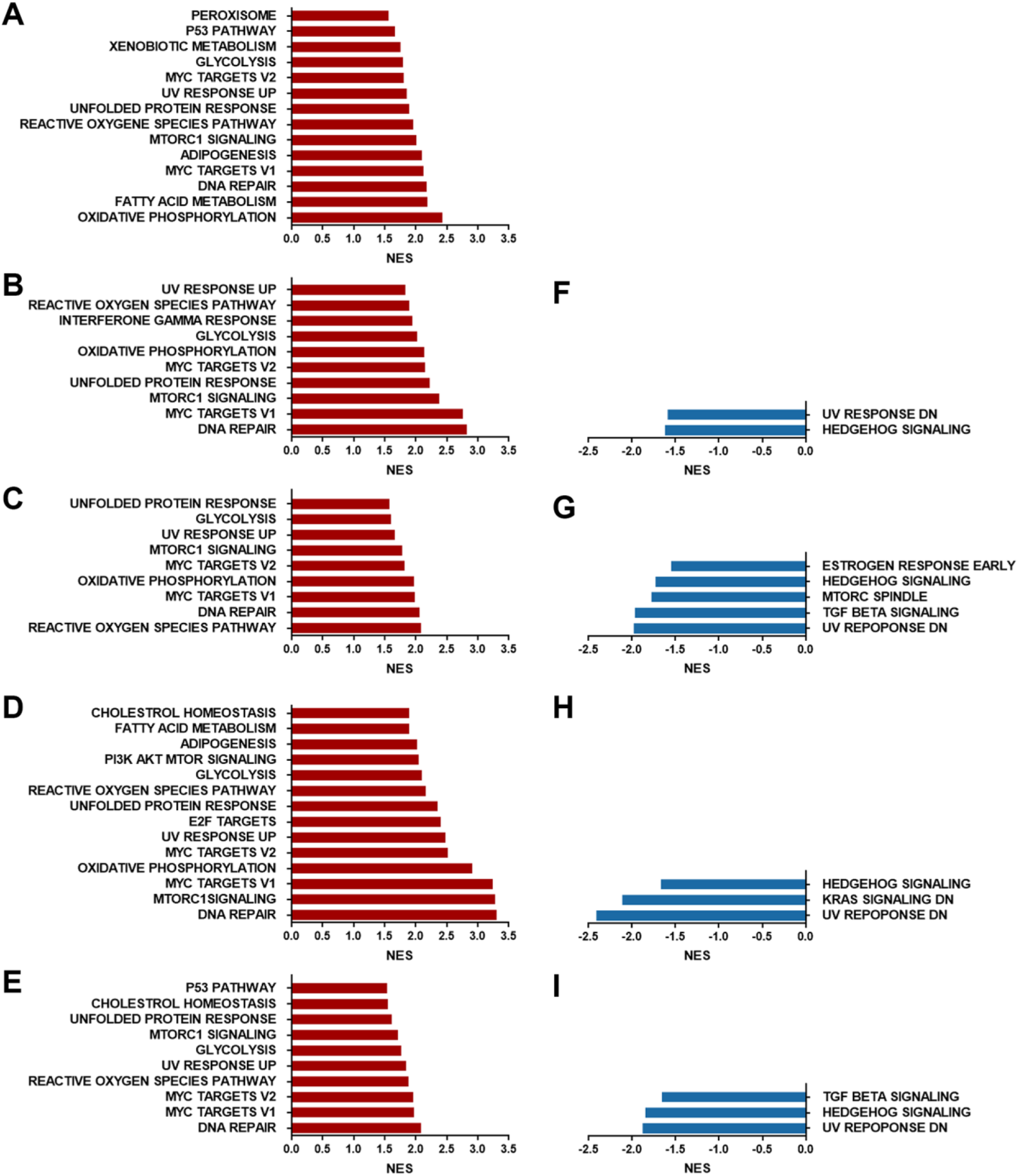
Pathway enrichment of 5 different cancer types given by GSEA analysis. Pathways enriched in samples with high expression of proteasome (A-E) or low expression of proteasome subunits (F-I) were shown in bar plots (in red for pathways enriched when proteasome is highly expressed and blue for pathways that associates with lower proteasome expression). Normalized Enriched scores (NES) was obtained from GSEA analysis. Significant enrichment was considered if p ≤ 0.05 and FDR ≤ 0.25. (A) data for LAML; (B and F) data for LGG; (C and G) data for LIHC; (D and H) data for LUAD and (E and I) data for KIRC.

### 3.5 Ingenuity pathway analysis of proteasome expression in cancer

The selected cancers KIRC, LAML, LGG, LIHC and LUAD were now classified based on changes in 14 proteasomal subunits and the proteasome maturation protein POMP using ingenuity pathway analysis (IPA). For this purpose, the intensities of gene expression of cancer type associated patients were used to divide patients into two cohorts (HIGH and LOW). Differentially expression values are available in the Supplementary Table S4. According to the classification into the respective cohorts, a comparison of gene expression between HIGH and LOW was performed for all genes included in the data set. A cut off of 1.5 was selected for the ingenuity pathway analysis (IPA). Supplementary Table S5 shows the numbers of differentially expressed genes used in the approach.

The most important pathway categories are cellular immune response, generation of precursor metabolites and energy, cellular growth, proliferation and development as well as neurotransmitter and other nervous system signaling (Table 1).

**Table 1:** z-scores for canonical pathways. The significant influenced pathways from IPA analysis were given by most increased or decreases z-scores.

| **category** | **KIRC** | **LAML** | **LGG** | **LIHC** | **LUAD** |
| --- | --- | --- | --- | --- | --- |
| cellular immune response | < -4 | < -3.5 | > 4.5 |  | < -4.5 |
| cellular stress and injury |  |  | > 4.5 |  |  |
| generation of precursor metabolites and energy | > 7 | > 5.5 |  | > 7 | > 7 |
| cellular growth, proliferation and development |  |  | < -8 | < -7 |  |
| neurotransmitter and other nervous system signaling |  |  | < -8 | < -7 |  |

The IPA analysis resulted in the canonical pathways shown in Figure 6 for the respective cancer type as bubble chart. For KIRC (Figure 6A) and LAML (Figure 6B) more of the genes showed an influence for activation of pathways, whereas in LGG, LIHC and LUAD (Figure 6C-E) more genes were annotated to inhibition of pathways. Additional information was given in the graphical summary of the analysis including entities such as canonical pathways, upstream regulators, diseases, and biological functions. The overview illustrates how those concepts relate to one another (comprehensive synopsis, Supplementary Figure S2).

**Figure 6:**
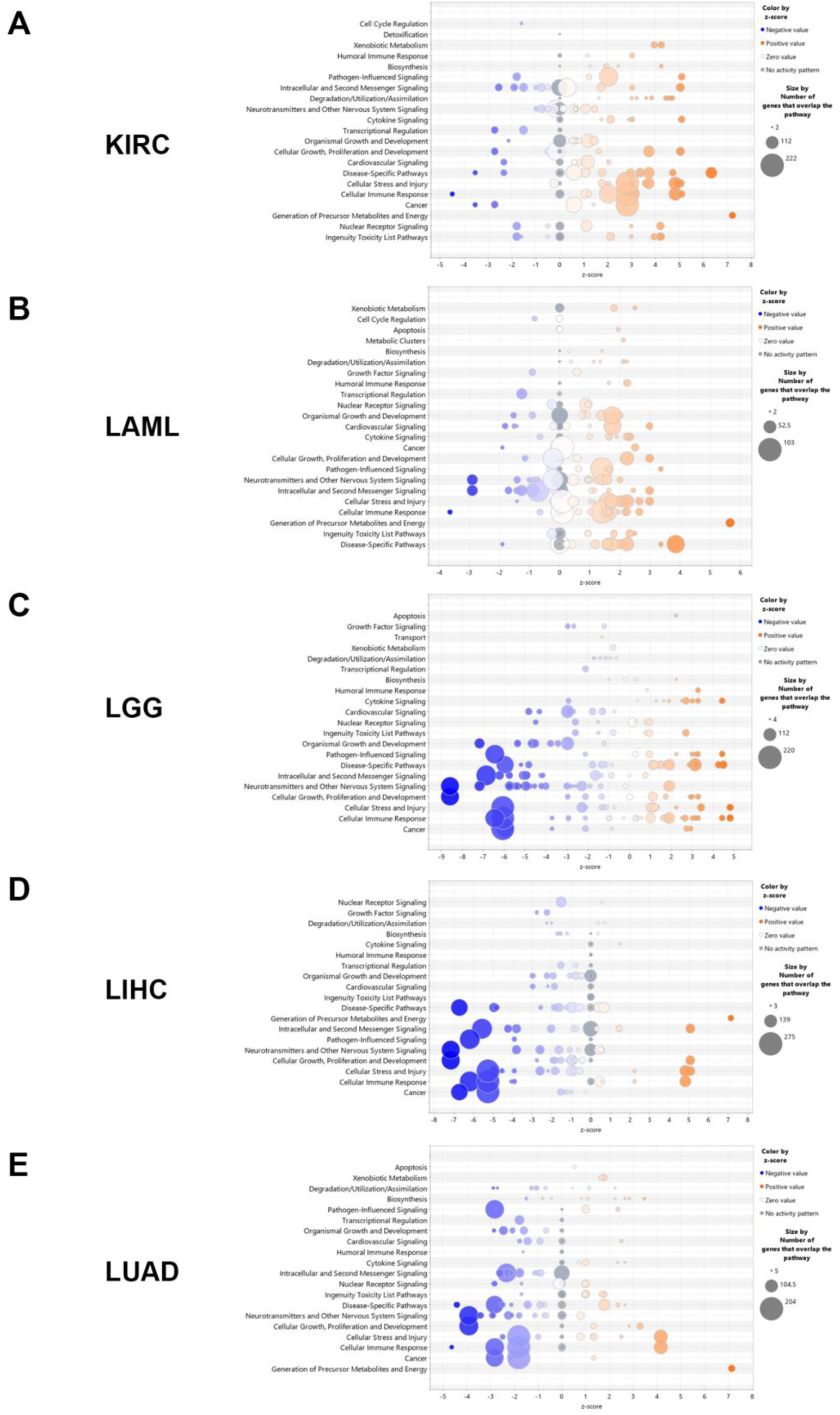
Canonical Pathways for 5 different cancer types given by the IPA analysis. The comparison of differentially expressed genes fitted to canonical pathway were shown as bubble chart plots; pathway categories (y-axis) versus the z-score (x-axis). The bubbles were coloured by z-score (blue-negative value; orange-positive value) and bubble size correlate to the number of analysis-ready genes from the dataset that overlap each pathway. (A) KIRC; (B) LAML; (C) LGG; (D) LIHC; (E) LUAD.

### 3.6 Pathway enrichment in THP-1 cells treated with proteasome inhibitor Bortezomib (BTZ)

As we have noticed that higher proteasome expression is associated with worse survival via the modification of oncogenic signaling pathways, we decided to confirm these changes on functional level in cells treated with the proteasome inhibitor Bortezomib (BTZ). For validation, we used THP-1 as a cancer cell model for LAML. After treatment with non-toxic dose of BTZ, RNA-seq followed by pathway enrichment analysis was carried out using GSEA. Proteasome inhibition caused enrichment in apoptosis pathway as well as p53 pathway. It also induced immune response related pathways such as NF-kB, interferon α response and inflammatory response (Figure 7). These pathways are known to sensitize cancer cells to immune therapy and enhance the immunogenic cell death (Figure 7A). As expected, on the metabolic level, while enrichment in the fatty acid metabolism and glycolysis was associated with high expression of proteasome subunits, we found that these pathways were downregulated after the proteasome inhibition using BTZ (Figure 7B). The most inhibited pathway was E2F targets which is known to play an essential role in cell proliferation and survival in cancer. Negative enrichment was observed in other oncogenic pathways such as MYC targets and oxidative phosphorylation (Figure 7C). Among the top 50 upregulated genes using proteasome inhibitor, genes involved in the innate immune signaling such as *ISG20* in addition to chemokines such as *CXCL3* and *CXCL8*. Other genes involved in fatty acid metabolism such as *PALM3*, *OLAH* beside a marked upregulation in the *ATF3* and the E3-ubiquitin ligase *TRIM54*. Furthermore, downregulation of genes from Wnt-β catenin signaling such as *WNT7B* and *CTNNA2* was observed after proteasome inhibition (Figure 7D). Indeed, our finding indicates that oncogenic and resistance pathways that were associated with high proteasome expression were downregulated when cancer cells are treated with proteasome inhibitors.

**Figure 7:**
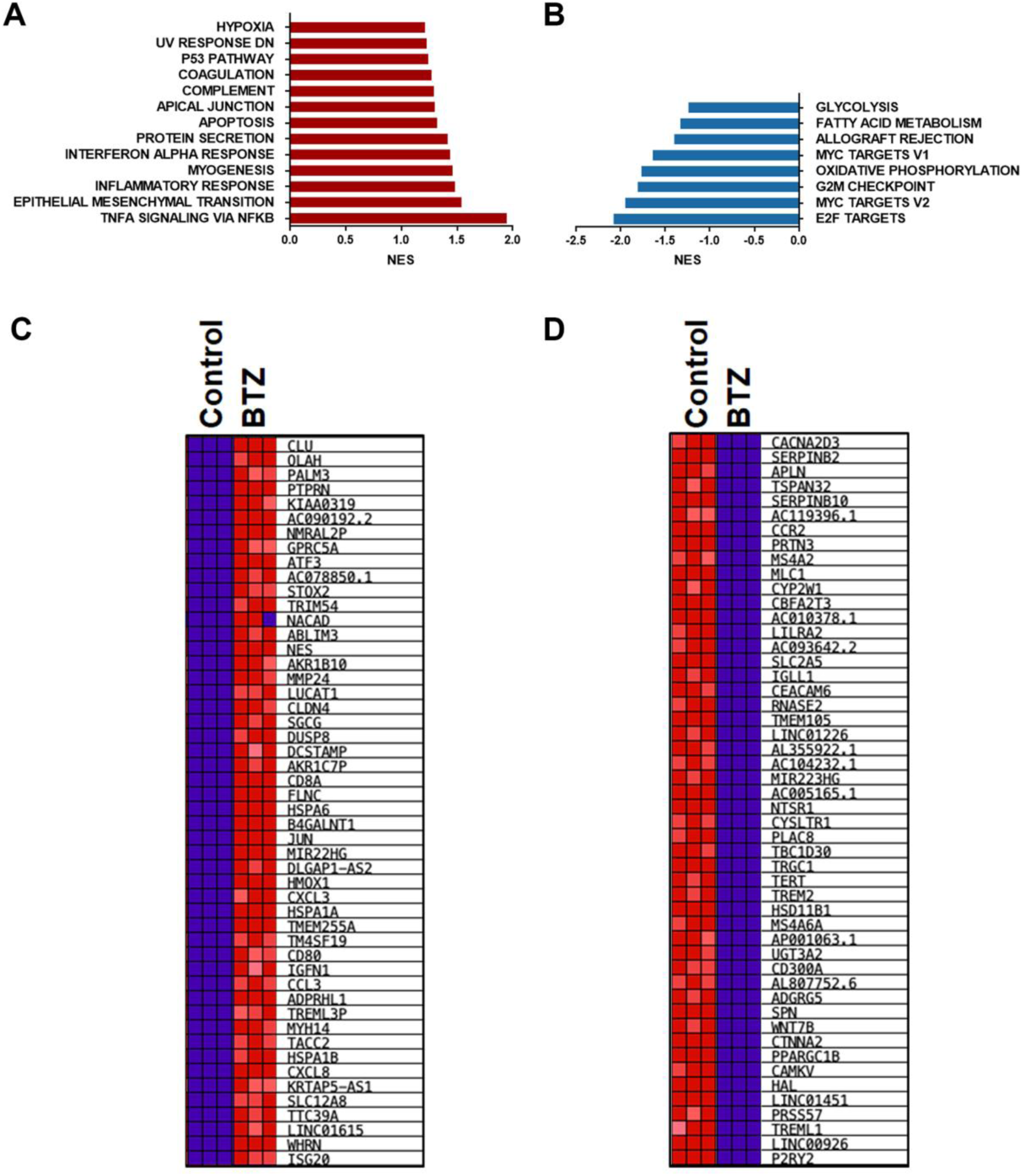
Pathway enrichment analysis and heatmap in cancer cells treated with proteasome inhibitor. RNA-seq followed by pathway analysis was carried for THP-1 cells after exposure to non-toxic dose (10 nM) of Bortezomib, BTZ for 16 hrs using GSEA analysis, results are presented as normalized enrichment score (NES). Pathway was considered enriched if p ≤ 0.05 and FDR ≤ 0.25. (A) enriched pathways in treated cells compared to the control; (B) enriched pathways in control compared to the treated cells; (C) Top 50 genes that shows the most elevated expression in treated cells.; (D) Top 50 genes that are downregulated genes in treated cells.

### 3.7 Correlation between gene expression of the proteasome 20S subunits and ABC transporters

ABC proteins are a family of transporters that participate in the reduction of intracellular concentrations of anti-tumor medications and therefore participate in the resistance developed against cancer treatment. However, the correlation of proteasome expression and ABC transporters is not fully studied. Among the most well-studied proteins from this family are *ABCG2* (*BCRP*), *ABCB1* (*MDR1*), and ABCC1 (*MRP1*). In this study, we created a signature of 24 transporters of the ABC proteins family (Supplementary Table S6). We used TCGA based-GEPIA2 database to create a correlation between the expression of proteasome subunits and ABC transporter. A positive significant correlation was observed in KIRC, LAML, LGG and, LIHC (Figure 8A-8D). However, this correlation was not observed in LUAD (Figure 8E). The effect of proteasome inhibition by Bortezomib on the expression of ABC transporters in THP-1 cells was variable (Figure 8F-G). The ABCCs subfamily of the efflux transporters is known to be involved in the resistance of conventional and adjacent cancer therapy [47]. Several genes involved in drug resistance such as *ABCC1* and *ABCC5* were slightly, however significantly, upregulated (Figure 8F) which could indicate that they could participate in the resistance to BTZ. Other ABC transporters such as *ABCC4* and *ABCC6* were downregulated after the treatment of these cells (Figure 8G). In this cell line, the expression levels of *ABCB1* and *ABCG2* were below the detection threshold.

**Figure 8:**
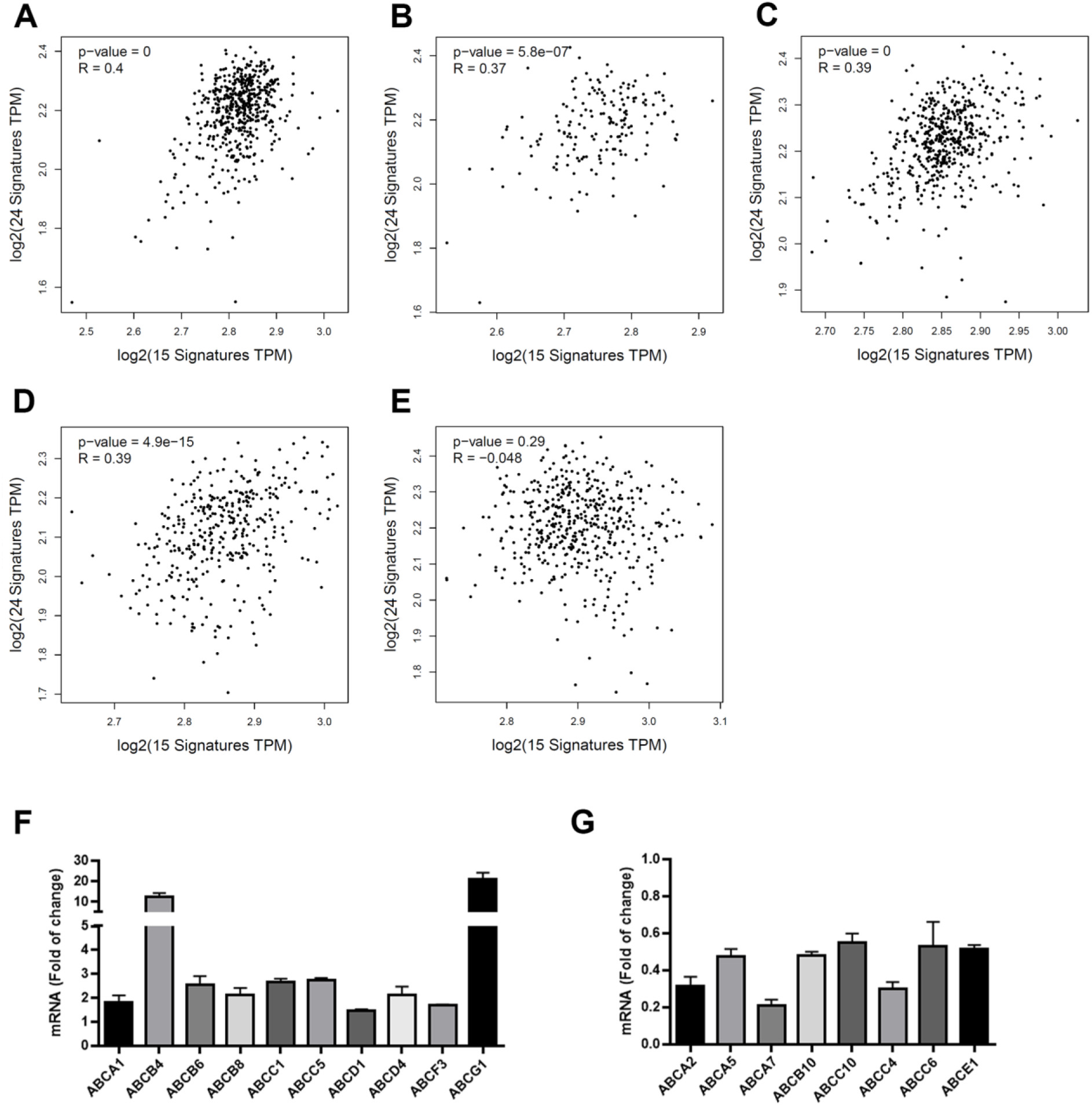
Correlation between proteasome 20S subunits expression and ABC transporters. (A - E) Correlation obtained using GEPIA2 database between the signature of proteasome (15 genes) and the signature of ABC transporters (24 genes). Results were considered significant if p≤0.05 and Pearson ≥ 0.3. (F and G) Transcriptome of ABC transporter in THP-1 cells after 16 hrs of exposure to 10 nM of Bortezomib. Results are shown as fold of changes of RPKM in treated cells when compared to control. Only significant results are shown ( p ≤0.05). Results are presented as mean ±SD.

## 4. Discussion

Proteasome inhibitors have emerged as novel therapeutic strategies in cancer management in the last two decades. These medications are currently the first-line treatments of hematopoietic malignancies including multiple myeloma and mantle cell lymphoma [48, 49]. The success of proteasome inhibitors in solid tumors is still limited and several clinical trials are still ongoing to estimate the effect in combination with other treatments. Therefore, it is essential to understand how proteasome expression is changed in cancer and which solid tumors could benefit from proteasome inhibition. In this study, we used the publicly available TCGA data to filter out tumor entities in which high expression of proteasome subunits is associated with worse patient survival. We have found that the effect of proteasome subunits expression on patient survival is cancer type-dependent. This phenomenon is predominant in hematopoietic malignancies such as multiple myeloma due to the elevated production of immunoglobins [50]. In LGG, we found a significant association between high transcriptome level of *PSMA6* and *PSMB5* and better prognosis. This effect is probably dependent on the cancer subtype. In ovarian cancer and to less degree in thyroid cancer, we found a better survival in patients with high proteasome expression. Indeed, a phase II-trial for ovarian cancer to establish the benefit of a combination of BTZ with doxorubicin was terminated because the anti-tumor activity failed [51]. However, other components of the UPS system like E3 ligases were elevated in epithelial type of ovarian cancer. These tumors could benefit from specific inhibitors of E3-ligases [52]. While the addition of low dose of decitabine, a DNA methylation inhibitor, could add to the effect of BTZ in the treatment of refractory multiple myeloma [53], the addition of bortezomib to decitabine did not improve the therapeutic outcome in acute myeloid leukemia patients [54]. Our results indicate that lower DNA methylation of the proteasome genes is often associated with worse survival, therefore we suggest that such combinations should be carefully designed. Our findings mainly indicate the importance of studying the proteasome as an enzymatic functional unit rather than separated subunits.

A previous analysis of PSMAs in breast cancer has indicated that except for *PSMA5* the expression of standard proteasome α-subunits was associated with worse survival [55]. Another study has pointed out the prognostic value of the β-subunits of the SP and IP in renal cell kidney cancer [25]. In contrast to these studies we evaluate the effect of the 20S proteasome as a functional unit including α- and β-subunits. We concentrated on cancer types, in which 20S proteasome subunit expression is consistently associated with worse survival.

DNA methylation has been found to have a significant impact on the expression of several proteasome subunits, particularly, in low grade glioma (LGG), and hepatocellular (LIHC). DNA methylation of *PSMA7* detected in the circulating DNA obtained from plasma of hepatocellular carcinoma patients could be used as a diagnosis and prognosis marker [56]. In breast cancer specific CpGs of *PSMA1*, *PSMA2*, *PSMA4*, *PSMA5*, *PSMA6* and *PSMA7* the highest levels of methylation were found [55]. It is worth mentioning that proteasome activity has an impact on epigenetic features including histone acetylation, the chromatin dynamics or stability, and DNA methylation [57]. The information is limited about the impact of DNA methylation on the expression of proteasome subunits in cancer and the prognostic and diagnostic value of them. In this analysis we identified several CpGs in LGG and LIHC that correlated negatively with gene expression of proteasome and could be candidates as prognostic markers.

While the variants identified in the TCGA somatic sequencing data have the potential to disrupt proteasome function, there is currently a lack of additional data to substantiate our variant predictions. Nonetheless, *PSMB4* has been identified as upregulated in various cancer types, including glioma [58–60], indicating that disruption of β7 encoded by *PSMB4* in TCGA-DU-6392 might itself contribute to impaired cancer cell metabolism. However, it is worth mentioning that an additive effect of multiple distinct subunit variants has been demonstrated to result in proteasomal loss of function and different diseases [14]. In this context, it is plausible to conclude that the collective presence of all the core particle variants found in sample TCGA-DU-6392 leads to proteasome impairment in the cancer cell, thereby contributing to the observed survival of this patient. Whether this effect occurs at a functional level or during proteasome assembly remains to be elucidated.

The UPS plays an essential role in various cellular functions including cell cycle, gene transcription, apoptosis, protein quality control, and epigenetic modifications [7, 57, 61].

The unfolded protein response pathway (UPR) was enriched in samples of patients with high proteasome subunit expression. In non-cancerous conditions elevated UPR is determined upon proteasome inhibition. The results observed here could be a direct result of the protein accumulation in cancer cells which was associated with worse survival and the enrichment of the UPR. Elevated expression of proteasome subunits might be a compensatory mechanism to cope with elevated proteotoxic stress. This phenomenon is also observed when cells were treated with proteasome inhibitors such as Bortezomib. The induced proteotoxic stress lead to the activation of TCF11/Nrf1 axis which results in elevated expression of proteasome subunits [37]. The higher expression of proteasome 20S subunits was combined with elevated cell survival pathways activation such as MYC targets, E2F targets, and DNA repair mechanisms. In contrast, the inhibition of the proteasome using BTZ leads to a significant reduction in the survival pathways mentioned and to the activation of apoptotic pathways in cancer cells. Metabolic pathways such as fatty acid metabolism, adipogenesis and glycolysis were enriched in the cohorts with high proteasome expression and reduced after treatment with BTZ. It has been reported that proteasome dysfunction disrupts adipogenesis and affects lipid metabolism [62].

There are controversial results about the interaction between proteasome inhibitors and ABC-family of transporters. For example, MG132, a proteasome inhibitor was found to reverse the resistance of vincristine-resistant human gastric cancer cell line by the inhibition of multi drug resistance MDR1 encoded by *ABCB1* [63]. In another study, it was demonstrated that BTZ treatment affected the expression of MRP1 encoded by *ABCC1* in a manner that depends on the AKT kinase activity [64]. In our cell model, we also observed a slight, but significant increase in MRP1 expression which might contribute to the resistance of tumors to BTZ in types of cancer with elevated *ABCC1* expression such as LIHC. On the other hand, further studies have reported the capacity of BTZ to reduce the expression of these resistance markers such as MDR1 and MRP1 [65], MDR1 via the inhibition of NF-kB pathway [66]. In myeloma patients treated with BTZ and doxorubicin, long term response was higher in patient with non-functional mutated MDR1 and MRP1 [67]. In our study, we have found that in THP-1 cells, BTZ did induce the expression of MRP1, which could contribute to BTZ resistance. However, we found a reduction in the expression of other efflux transporters such as MRP4. These results suggest that BTZ could reduce the resistance to drugs that are substrates for these transporters such as 5-Fluorouracil.

## 5. Conclusion

In general, our study pointed out that, albeit the effect that is observed for specific proteasome subunits on cellular function, the main role of proteasome is better evaluated when the proteasome is studied as an enzyme unit. In this comprehensive study, we have defined cancer types in which proteasome subunits expression is associated with worse survival. These types of cancer are candidates to develop therapeutic strategies that aim to inhibit the proteasomal activity or induce proteo-toxic stress. We also detected DNA methylation sites that are associated with reduced expression. In addition to the well-defined pathways that could be affected by proteasome expression, we explored novel correlation between proteasome subunit expression, the expression of ABC transporters, and stem cells markers in various types of cancer.

## Data accessibility

The FastQ files of the RNA-seq experiment is shared in GEO under PRJNA1032858.

## Authors contributions

Manuscript Design (data and experimental design): EK and RAA Data processing: RAA, SV, RA

Mutations data: MW, SV and RAA Manuscript preparation: RAA, SV, and EK RNA-seq experiment: RAA, MR

RNA-seq analysis: RAA and RA Statistical analysis: RAA, EK and SV

## Acknowledgements

We thank the Interfaculty Institute of Genetics and Functional Genomics, University Medicine Greifswald for support with the Ingenuity Pathway Analysis (IPA). We thank Dr. Heike Junker for her help with the RNA extraction protocol.

## Abbreviations

BTZ: Bortezomib
UPS: Ubiquitin Proteasome System
UPR: Unfolded Protein Response pathway
CP: core particle
RP: regulatory particle
SP: standard proteasome
IP: immunoproteasome
GSEA: Gene Set Enrichment Analysis
ACMG: The American College of Medical Genetics and Genomics
IPA: Ingenuity Pathway Analysis
QC: Quality Control
NES: Normalized Enrichment Score
TCGA: The Cancer Genome Atlas.

## Supporting Information

Supplementary Table S1:

TCGA abbreviations and number of samples used in the survival study.

Supplementary Table S2:

Proteins encoded by the proteasome subunits genes.

Supplementary Table S3:

Identification of potentially disruptive variants in proteasome-associated genes.

Supplementary Table S4:

Differential expression factors of gene of interest by mean values.

Supplementary Table S5:

Number of classified patients out of the whole cancer specific patient number.

Supplementary Table S6:

24 transporters of the ABC proteins family.

Supplementary Figure S1:

Electrostatic potential and surface hydrophobicity of the variants discovered in TCGA-DU-6392, visualized in ChimeraX.

Supplementary Figure S2:

Graphical summary from the IPA core analysis.

