## Supplementary material for "Expression and DNA methylation of 20S proteasome subunits as prognostic and resistance markers in cancer": suplementary data

Supplementary Table S1:
TCGA abbreviations and number of samples used in the survival study.

Supplementary Table S2:
Proteins encoded by the proteasome subunits genes.

Supplementary Table S3:
Identification of potentially disruptive variants in proteasome-associated genes.

Supplementary Table S4:
Differential expression factors of gene of interest by mean values.

Supplementary Table S5:
Number of classified patients out of the whole cancer specific patient number.

Supplementary Table S6:
24 transporters of the ABC proteins family.

Supplementary Figure S1:
Electrostatic potential and surface hydrophobicity of the variants discovered in TCGA-DU-6392, visualized in ChimeraX.

Supplementary Figure S2:
Graphical summary from the IPA core analysis.

**Supplementary Table S1:**

**TCGA abbreviations and number of samples used in the survival study.**

| Abbreviation | Cancer type | Nr. of Patients |
|---|---|---|
| LAML | Acute Myeloid Leukemia | 151 |
| LGG | Low Grade Glioma | 511 |
| KIRC | Kidney Renal Cell Carcinoma | 538 |
| LIHC | Liver Hepatocellular Carcinoma | 371 |
| LUAD | Lung Adenocarcinoma | 533 |
| OV | Ovarian Serous Cystadenocarcinoma | 374 |
| THYM | Thymoma | 119 |
| THCA | Thyroid Carcinoma | 502 |
| ESCA | Esophageal Carcinoma | 161 |
| HNSC | Head and Neck Squamous Cell Carcinoma | 500 |
| KIRP | Kidney Renal papillary Cell Carcinoma | 288 |
| LUSC | Lung Squamous Cell Carcinoma | 502 |
| PAAD | Pancreatic Adenocarcinoma | 177 |
| PRAD | Prostate Adenocarcinoma | 498 |
| SARC | Sarcoma | 259 |
| STAD | Stomach Adenocarcinoma | 375 |
| TGCT | Testicular Germ Cell Tumors | 150 |
| UCEC | Uterine Corpus Endometrial Carcinoma | 551 |
| BLCA | Bladder Urothelial Carcinoma | 414 |
| GBM | Glioblastoma Multiforme | 156 |
| BRCA | Breast Invasive Carcinoma | 1102 |
| CESC | Cervical Squamous Cell Carcinoma and Endocervical Adenocarcinoma | 304 |
| COAD | Colon Adenocarcinoma | 478 |
| READ | Rectum Adenocarcinoma | 304 |

**Supplementary Table S2:**

**Proteins encoded by the proteasome subunits genes.**

| Proteasome alpha subunits | Proteasome beta subunits | ATPase – dependent regulatory particles (RPTs) | ATPase independent regulatory genes | Other Regulators |
|---|---|---|---|---|
| *PSMA1* (α6 subunit) | *PSMB1* (β6 subunit) | *PSMC1* (Rpt2) | *PSMD1* (Rpn2) | *PSME1* (PA28α) |
| *PSMA2* (α2 subunit) | *PSMB2* (β4 subunit) | *PSMC2* (Rpt1) | *PSMD2* (Rpn1) | *PSME2* (PA28β) |
| *PSMA3* (α7 subunit) | *PSMB3* (β3 subunit) | *PSMC3* (Rpt5) | *PSMD3* (Rpn3) | *PSME3* (PA28γ) |
| *PSMA4* (α3 subunit) | *PSMB4* (β7 subunit) | *PSMC4* (Rpt3) | *PSMD4* (Rpn10) | *PSME4* (PA200) |
| *PSMA5*(α5 subunit) | *PSMB5* (β5 subunit) | *PSMC5* (Rpt6) | *PSMD6* (Rpn7) | *PSMF1* (PI31) |
| *PSMA6* (α1 subunit) | *PSMB6* (β1 subunit) | *PSMC6* (Rpt4) | *PSMD7* (Rpn8) | |
| *PSMA7* (α4 subunit) | *PSMB7* (β2 subunit) | | *PSMD8* (Rpn12) | |
| *PSMA8* (α7-like protein) | *PSMB8* (β5i subunit) | | *PSMD11* (Rpn6) | |
| | *PSMB9* (β1i subunit) | | *PSMD12* (Rpn5) | |
| | *PSMB10* (β2i, MECL1 subunit) | | *PSMD13* (Rpn9) | |
| | *PSMB11* (β5t subunit) | | *PSMD14* (Rpn11) | |
| | | | *ADRM1* (Rpn13) | |
| | | | *SHFM1* (Rpn15) | |

**Supplementary Table S3:**

**Identification of potentially disruptive variants in proteasome-associated genes.**
extraction, filtering, and manual classification of TCGA somatic sequencing data

| Cancer type | TCGA Tumor Sample | Genes | HGVS | ACMG Class |
|---|---|---|---|---|
| KIRC | TCGA-B8-5164 | POMP | c.162+1G>T | 4 |
| | TCGA-BP-4964 | PSMA3 | c.396dup p.(Gly133Trpfs*13) | 4 |
| | TCGA-B0-4823 | PSMA4 | c.581delinsGCAAAG p.(Ile194Serfs*6) | 4 |
| | TCGA-B0-4811 | PSMA6 | c.77-1G>C | 4 |
| | TCGA-B0-5088 | PSMB4 | c.374_375insT p.(His126Thrfs*4) | 4 |
| LAML | - | - | - | - |
| LGG | TCGA-P5-A77X | PSMA2 | c.226T>C p.(Tyr76His) | 3 |
| | TCGA-DU-6392 | PSMB4 | c.257G>A p.(Arg86His) | 3 |
| LIHC | TCGA-G3-A3CK | PSMA4 | c.356A>T p.(Gln119Leu) | 3 |
| | TCGA-ED-A7PZ | PSMA7 | c.527A>G p.(Tyr176Cys) | 3 |
| | TCGA-2Y-A9H0 | PSMB2 | c.317G>A p.(Gly106Asp) | 3 |
| | TCGA-WQ-A9G7 | PSMB3 | c.296+2T>C | 3 |
| LUAD | TCGA-MN-A4N5 | PSMA1 | c.238G>T p.(Asp80Tyr) | 3 |
| | TCGA-05-4427 | PSMA3 | c.524C>T p.(Thr175Met) | 3 |
| | TCGA-50-6590 | PSMA6 | c.245G>C p.(Gly82Ala) | 3 |
| | TCGA-55-8203 | PSMA7 | c.348+1G>A | 4 |
| | TCGA-78-8660 | | c.389C>T p.(Ser130Phe) | 3 |

**Supplementary Table S4:**

**Differential expression factors of gene of interest by mean values.**
Fold change factor was calculated by the gene expression data of all patients fitting to cancer type. The significance is given by the determined p-value.

| **Hugo Symbol** | **KIRC factor** | KIRC pvalue | **LAML factor** | LAML pvalue | **LGG factor** | LGG pvalue | **LIHC factor** | LIHC pvalue | **LUAD factor** | LUAD pvalue |
|---|---|---|---|---|---|---|---|---|---|---|
| ***PSMA1*** | **1.29** | 2.16E-30 | **1.19** | 7.53E-09 | **1.19** | 1.33E-25 | **1.31** | 2.83E-21 | **1.43** | 1.25E-32 |
| ***PSMA2*** | **1.32** | 3.38E-22 | **1.37** | 1.77E-15 | **1.32** | 1.67E-42 | **1.35** | 6.49E-16 | **1.59** | 1.94E-41 |
| ***PSMA3*** | **1.45** | 2.33E-36 | **1.32** | 3.55E-15 | **1.31** | 3.98E-39 | **1.48** | 1.11E-30 | **1.64** | 1.58E-32 |
| ***PSMA4*** | **1.46** | 2.39E-53 | **1.25** | 1.25E-09 | **1.30** | 1.37E-31 | **1.56** | 9.72E-38 | **1.54** | 3.25E-39 |
| ***PSMA5*** | **1.51** | 1.05E-47 | **1.42** | 9.44E-22 | **1.34** | 7.87E-34 | **1.61** | 7.66E-26 | **1.68** | 2.05E-48 |
| ***PSMA6*** | **1.32** | 5.50E-10 | **1.28** | 6.48E-12 | **1.28** | 9.80E-35 | **1.52** | 6.07E-35 | **1.82** | 1.06E-15 |
| ***PSMA7*** | **1.49** | 2.70E-18 | **1.46** | 3.14E-19 | **1.39** | 1.35E-50 | **1.68** | 2.19E-35 | **1.62** | 6.49E-33 |
| ***PSMB1*** | **1.49** | 5.51E-47 | **1.32** | 1.03E-24 | **1.36** | 1.93E-36 | **1.94** | 9.29E-36 | **1.66** | 6.69E-48 |
| ***PSMB2*** | **1.32** | 2.35E-34 | **1.10** | 3.75E-03 | **1.20** | 5.97E-18 | **1.49** | 4.61E-25 | **1.42** | 3.48E-36 |
| ***PSMB3*** | **1.97** | 8.75E-42 | **1.48** | 1.85E-15 | **1.56** | 5.99E-45 | **2.13** | 2.95E-32 | **1.79** | 4.94E-32 |
| ***PSMB4*** | **1.63** | 4.25E-53 | **1.36** | 2.30E-16 | **1.37** | 7.69E-48 | **1.68** | 4.37E-27 | **1.58** | 1.93E-33 |
| ***PSMB5*** | **1.35** | 1.03E-26 | **1.42** | 3.90E-14 | **1.23** | 1.52E-21 | **1.39** | 1.99E-24 | **1.67** | 3.09E-32 |
| ***PSMB6*** | **1.63** | 5.09E-44 | **1.48** | 1.58E-21 | **1.40** | 4.37E-43 | **1.85** | 1.80E-27 | **1.64** | 2.75E-37 |
| ***PSMB7*** | **1.49** | 4.11E-41 | **1.31** | 5.56E-17 | **1.31** | 3.96E-29 | **1.84** | 2.33E-35 | **1.63** | 1.14E-44 |
| ***POMP*** | **1.46** | 1.91E-54 | **1.51** | 1.34E-15 | **1.30** | 2.37E-31 | **1.75** | 1.80E-37 | **1.68** | 5.68E-24 |

**Supplementary Table S5:**

**Number of classified patients out of the whole cancer specific patient number.** To pass the IPA criteria for the pathway analysis a cut off factor of 1.5 fold was used to diminish the number of different genes within the cancer types.

| Disease | No. of expressed genes (TCGA) | No. of differentially expressed genes factor 1.5 |
|---|---|---|
| **KIRC** | 17637 | 4925 |
| **LAML** | 17729 | 2119 |
| **LGG** | 17607 | 3790 |
| **LIHC** | 17499 | 5834 |
| **LUAD** | 17582 | 4534 |

**Supplementary Table S6:**

**24 transporters of the ABC proteins family.** These proteins were used to create a correlation between the expression of proteasome subunits and ABC transporter.

| Gene Names (primary) | Entry | Entry Name (HUMAN) | Protein names |
|---|---|---|---|
| ABCA1 | O95477 | ABCA1 | Phospholipid-transporting ATPase ABCA1 |
| ABCA2 | Q9BZC7 | ABCA2 | ATP-binding cassette sub-family A member 2 |
| ABCA3 | Q99758 | ABCA3 | Phospholipid-transporting ATPase ABCA3 |
| ABCA5 | Q8WWZ7 | ABCA5 | Cholesterol transporter ABCA5 |
| ABCA7 | Q8IZY2 | ABCA7 | Phospholipid-transporting ATPase ABCA7 |
| ABCB10 | Q9NRK6 | ABCBA | ATP-binding cassette sub-family B member 10, mitochondrial |
| ABCB4 | P21439 | MDR3 | Phosphatidylcholine translocator ABCB4 |
| ABCB6 | Q9NP58 | ABCB6 | ATP-binding cassette sub-family B member 6 |
| ABCB7 | O75027 | ABCB7 | Iron-sulfur clusters transporter ABCB7, mitochondrial |
| ABCB8 | Q9NUT2 | MITOS | Mitochondrial potassium channel ATP-binding subunit |
| ABCB9 | Q9NP78 | ABCB9 | ABC-type oligopeptide transporter ABCB9 |
| ABCC1 | P33527 | MRP1 | Multidrug resistance-associated protein 1 |
| ABCC10 | Q5T3U5 | MRP7 | ATP-binding cassette sub-family C member 10 |
| ABCC4 | O15439 | MRP4 | ATP-binding cassette sub-family C member 4 |
| ABCC5 | O15440 | MRP5 | ATP-binding cassette sub-family C member 5 |
| ABCC6 | O95255 | MRP6 | ATP-binding cassette sub-family C member 6 |
| ABCD1 | P33897 | ABCD1 | ATP-binding cassette sub-family D member 1 |
| ABCD3 | P28288 | ABCD3 | ATP-binding cassette sub-family D member 3 |
| ABCD4 | O14678 | ABCD4 | Lysosomal cobalamin transporter ABCD4 |
| ABCE1 | P61221 | ABCE1 | ATP-binding cassette sub-family E member 1 |
| ABCF1 | Q8NE71 | ABCF1 | ATP-binding cassette sub-family F member 1 |
| ABCF2 | Q9UG63 | ABCF2 | ATP-binding cassette sub-family F member 2 |
| ABCF3 | Q9NUQ8 | ABCF3 | ATP-binding cassette sub-family F member 3 |
| ABCG1 | P45844 | ABCG1 | ATP-binding cassette sub-family G member 1 |

**Supplementary Figure S1:**

**Electrostatic potential and surface hydrophobicity of the variants discovered in TCGA-DU-6392, visualized in ChimeraX.** (A-F) Comparison of the electrostatic potential (top panel) and hydrophobicity (bottom panel) between the reference structure of each subunit (left) and the corresponding folded variant structure (right).

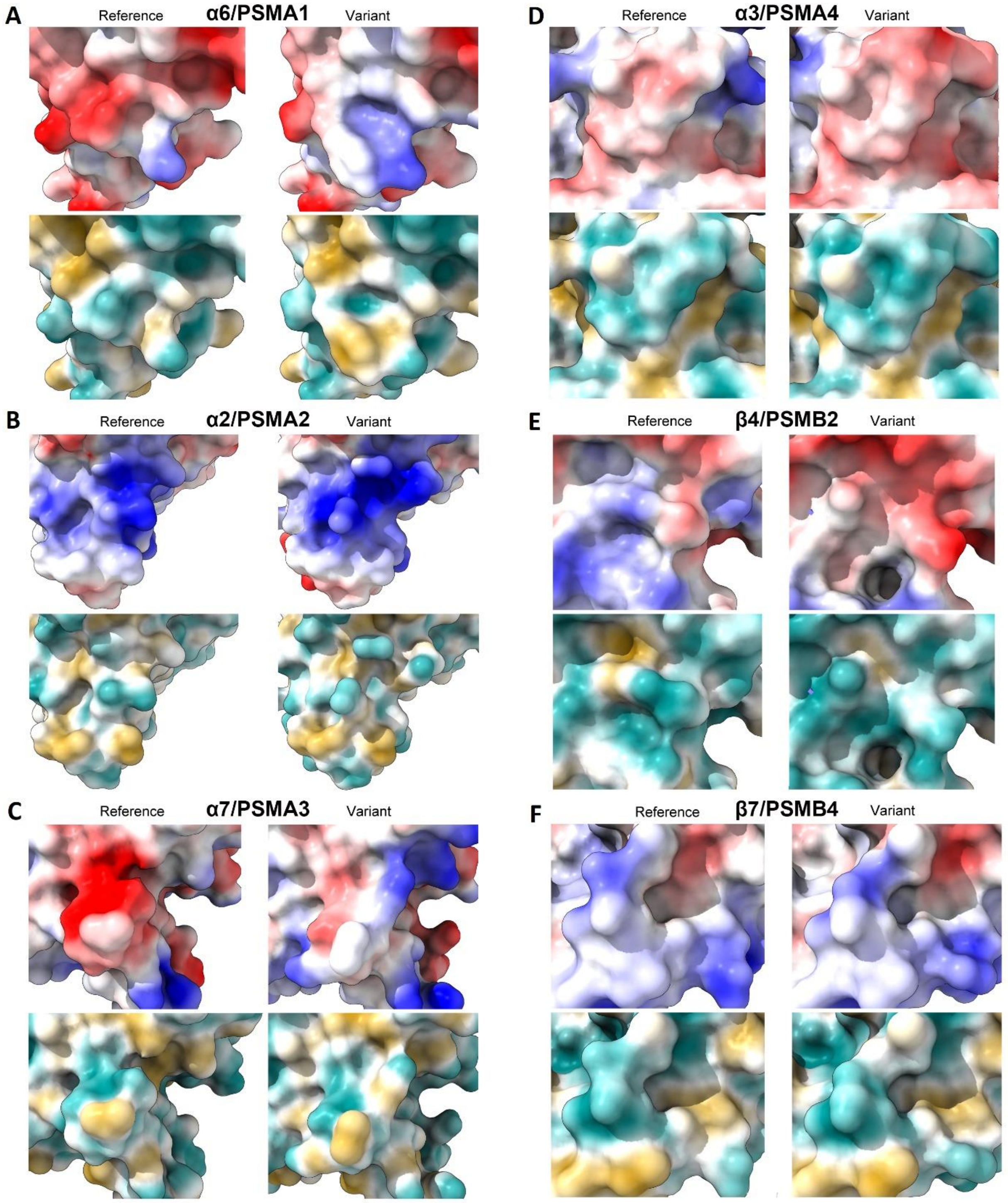

**Supplementary Figure S2:**

**Graphical summary from the IPA core analysis.**

Summary of key results from the IPA core analysis for each type of cancer presented in a network. Predicted activation were coloured with orange symbols and lines. Predicted inactivation were coloured with blue symbols and lines. Marks in grey fit to interaction behavior.

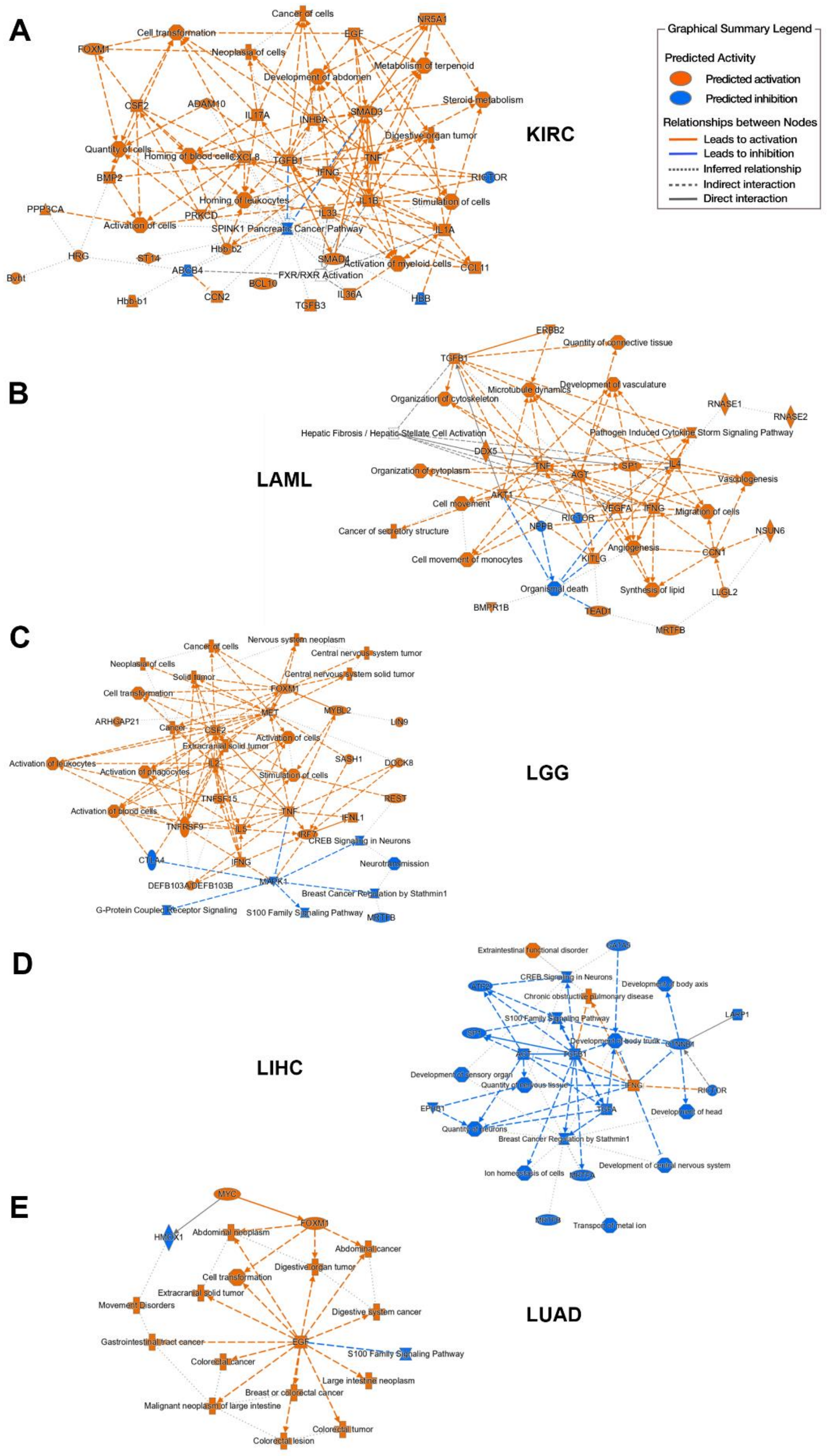
